# Redox regulation of PTPN22 affects the severity of T cell dependent autoimmune inflammation

**DOI:** 10.1101/2021.12.02.470976

**Authors:** Jaime James, Yifei Chen, Clara M. Hernandez, Florian Forster, Markus Dagnell, Qing Cheng, Amir A. Saei, Hassan Gharibi, Gonzalo Fernandez Lahore, Annika Åstrand, Rajneesh Malhotra, Bernard Malissen, Roman A. Zubarev, Elias S.J. Arnér, Rikard Holmdahl

**Author notes:** Corresponding author, Rikard Holmdahl, ^1^Division of Medical Inflammation Research, Dept. of Medical Biochemistry and Biophysics, Karolinska Institute, Stockholm, Sweden. **Classification:** Biological Sciences/Immunology and Inflammation.

## Abstract

Chronic autoimmune diseases are associated with mutations in PTPN22, a modifier of T cell receptor signaling. As with all protein tyrosine phosphatases the activity of PTPN22 is redox regulated, but if or how such regulation can modulate inflammatory pathways *in vivo* is not known. To determine this, we created a mouse with a cysteine-to-serine mutation at position 129 in PTPN22 (C129S), a residue proposed to alter the redox regulatory properties of PTPN22 by forming a disulfide with the catalytic C227 residue. The C129S mutant mouse showed a stronger T cell-dependent inflammatory response and development of T cell dependent autoimmune arthritis due to enhanced TCR signaling and activation of T cells, an effect neutralized by a mutation in Ncf1, a component of the NOX2 complex. Activity assays with purified proteins suggest that the functional results can be explained by an increased sensitivity to oxidation of the C129S mutated PTPN22 protein. We also observed that the disulfide of native PTPN22 can be directly reduced by the thioredoxin system, while the C129S mutant lacking this disulfide was less amenable to reductive reactivation. In conclusion, we show that PTPN22 functionally interacts with Ncf1 and is regulated by oxidation via the non-catalytic C129 residue and oxidation-prone PTPN22 leads to increased severity in the development of T cell-dependent autoimmunity.

**Significance statement:** A hitherto unstudied aspect of PTPN22 biology is its regulation by cell redox states. Here we created a mouse model where PTPN22 was mutated to respond differentially to redox levels *in vivo* and found that PTPN22 function is regulated by reactive oxygen species and that redox regulation of PTPN22 impacts T-cell-dependent autoimmune inflammation.

## Introduction

Complex autoimmune diseases affect 4-5% of the human population and large efforts have been invested in finding the underlying genetic polymorphisms [1]. Though a major genetic contribution comes from the major histocompatibility complex region, many other loci have also been identified. Two important single nucleotide polymorphisms (SNPs) have emerged, one located in PTPN22 [2], a cytoplasmic class I protein tyrosine phosphatase (PTP), and the other in NCF1 [3][4], a component of the NOX2 complex, controlling induction of reactive oxygen species (ROS) in antigen presenting cells. Mutations in *Ncf1*, leading to a lower NOX2 dependent ROS response, have been shown to be a major predisposing genetic factor for autoimmune diseases in both mice and humans [3][4]. PTPN22 is primarily a dominant negative regulator of T cell responsiveness, acting by dephosphorylating important target proteins in the T cell signaling machinery including LCK, FYN and ZAP70 [5][6]. Studies on PTPN22 have thus far focused on the effects of knocking out the gene or on the autoimmune variant *PTPN22*-*R620W*, but with divergent results [7]. Importantly, however, PTPN22 is likely to be redox regulated. Its activity is, as with other protein tyrosine phosphatases (PTPs), dependent upon the integrity of a catalytic active site Cys residue. The reactivity of this cysteine makes it particularly susceptible to oxidation via reactive oxygen species (ROS) leading to concomitant abrogation of PTP activity. Reversible oxidation and reduction of such reactive cysteines has previously been shown to be a regulatory mechanism in signal transduction, regulating important T cell downstream signaling molecules such as NF-KB, NRF2, MAPK and ERK [8]-[9]. Control of redox regulation of PTPs in cells depends upon the balance between inhibitory oxidation of the catalytic Cys residue and its activation by reduction, where the latter is typically maintained by the thioredoxin system. We have now investigated the possibility that redox regulation of PTPN22 could modulate inflammatory pathways *in vivo.* Interestingly, the PTPN22 catalytic cysteine (C227) has been suggested to form a disulfide bond with another “back-door” cysteine (C129), possibly altering the threshold for irreversible oxidation of C227 and thereby affecting the redox state of the active site [10]. By creating a mouse with a cysteine-to-serine mutation at position 129 in PTPN22 (C129S) we could investigate the possible pathophysiological impact of its redox regulatory properties *in vivo*. We found that mice with this amino acid replacement developed increased T cell-dependent inflammatory responses due to enhanced T cell receptor signaling, which was dependent on NOX2-produced ROS. This correlated well with findings of a lower overall turnover, higher sensitivity to inhibitory oxidation, and a lower capacity of reductive reactivation by the thioredoxin system of recombinant mutant PTPN22^C129S^, as compared to wild-type PTPN22. These results show that redox regulation of PTPN22 modulates inflammation *in vivo*, with a lower resistance to oxidation of PTPN22 promoting aggravated disease.

## Results

### Recombinant PTPN22^C129S^ has lower catalytic activity, higher sensitivity to inhibition by oxidation and lower capacity for reductive reactivation by the thioredoxin system

To investigate the potential impact of a C129S replacement in the PTPN22 protein, we recombinantly produced the catalytic domain of the corresponding human wild-type PTPN22 and PTPN22^C129S^ mutant proteins (the C129 and the catalytic C227 residues have the same numbering in both mouse and human). The recombinant proteins were purified to >95% purity as judged by SDS-PAGE and kinetic parameters were determined using p-NPP as substrate; wild-type PTPN22 enzyme had a basal turnover of 19.6 min^-1^ with a K_m_ of 4.57 mM while PTPN22^C129S^ displayed a turnover of 11.9 min^-1^ and a K_m_ of 8.02 mM under the same conditions, showing that the C129S protein has retained PTP activity but with an overall lower catalytic efficiency (Fig. 1A) which confirms Tsai et al.’s results [10].

**Fig.1.**
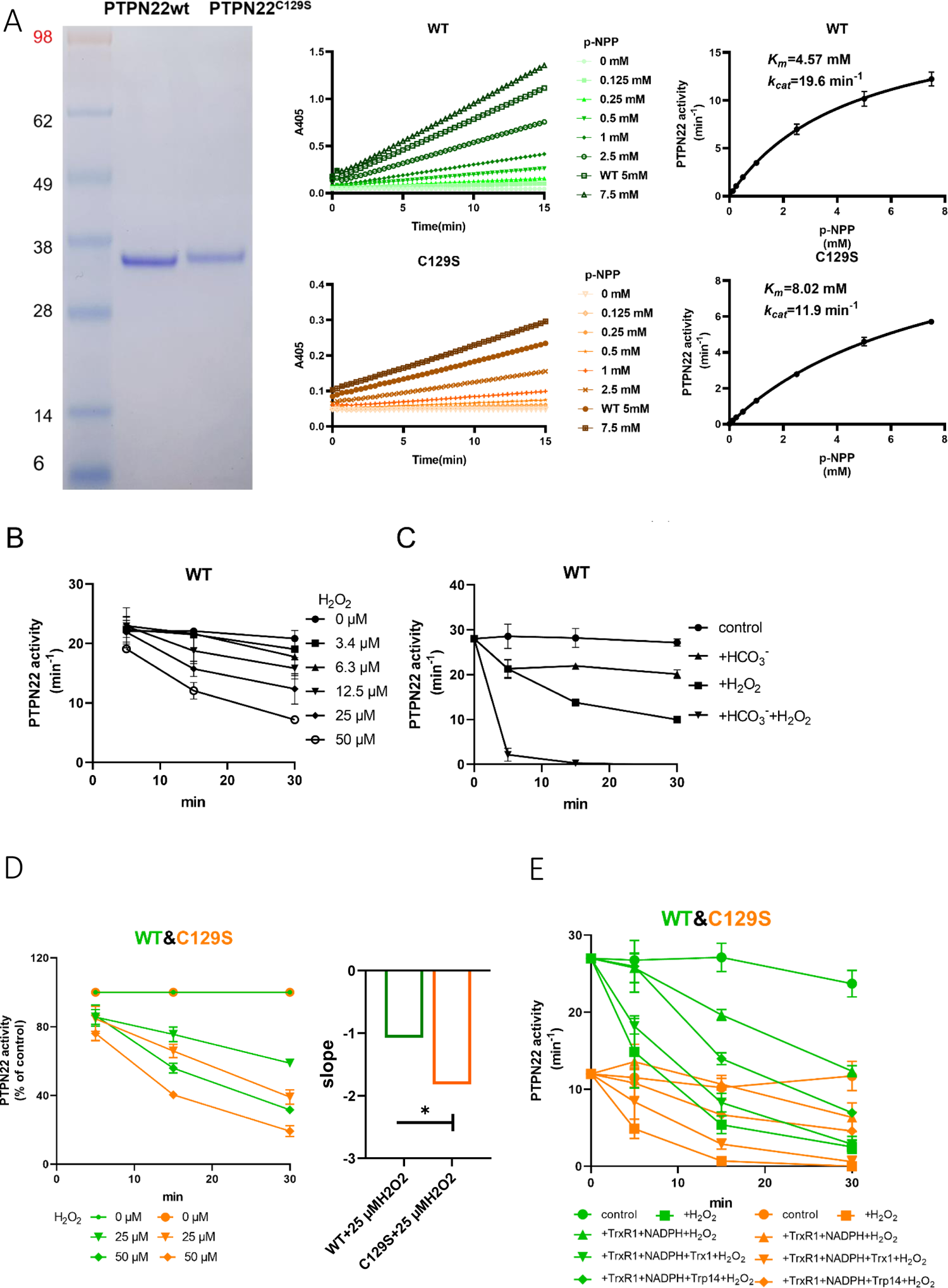
PTPN22C129S is more prone to oxidative inactivation. **A)** Purified human wild-type and PTPN22^C129S^ proteins are shown as analyzed on SDS-PAGE, and kinetic parameters were measured using p-NPP as substrate as shown to the right. Data points represent mean ± S.D. (error bars) (n=3). **B)** PTPN22 (1400nM) was treated with indicated concentrations of H_2_O_2_ and then assayed for PTP activity at the indicated times. PTPN22 activity is given in min^-1^ (mol of product/mol of enzyme/min). Data points represent mean ± S.D. (error bars) (n=3). **C)** PTPN22 (1400nM) activity was measured as in *A* but with buffer control (●), 25mM bicarbonate (▲), 30µM H_2_O_2_ (▪) or the combination of 30µM H_2_O_2_ and 25mM bicarbonate (▼). Data points represent mean ±S.D. (error bars) (n=4). **D)** Wild-type (green) and PTPN22^C129S^ (orange) proteins were treated with indicated H_2_O_2_ concentrations and then assayed for PTP activity. Data points represent mean ±S.D. (error bars) (n=3) and activities are given as % of buffer control for each enzyme. **E)** Pre-reduced wild-type (green) and PTPN22^C129S^ (orange) proteins were first preincubated for 10 min either in only buffer (▪), with 300μM NADPH and 0.25μM TrxR1 (▲), with 300μM NADPH, 0.25μM TrxR1 and 10μM Trx1(▼), or with 300μM NADPH, 0.25μM TrxR1 and 10μM Trp14(◆), whereupon H_2_O_2_ (100μM) and 1mM NaN_3_ was added, except to the control (●), and samples were taken at indicated times for measurement of PTP activity. Data points represent mean ±S.D. (error bars) (n=3).

Next, we wanted to assess sensitivity of pre-reduced wild-type PTPN22 to inhibition by oxidation. We found that addition of 50 µM H_2_O_2_ to the pure protein led to inhibition of approximately half the activity after 20 min incubation (Fig. 1B). We also found that addition of bicarbonate together with H_2_O_2_ noticeably potentiated the inactivation (Fig. 1C), similar to the properties shown earlier for PTP1B that are likely due to formation of peroxymonocarbonate that reacts more efficiently than H_2_O_2_ with the PTP enzyme [11]. When comparing the H_2_O_2_-sensitivity of PTPN22^C129S^ with that of wild-type PTPN22, we found that the mutant enzyme was clearly more sensitive to inhibition by H_2_O_2_ than wild-type PTPN22, although it had a lower basal turnover (Fig. 1D). The same effect was also seen in the presence of a functional thioredoxin system of thioredoxin reductase 1 (TrxR1) coupled with thioredoxin (Trx1), but perhaps less so coupled with thioredoxin related protein of 14 kDa (TRP14) (Fig. 1E). It should be noted that also PTP1B displays different reactivities with Trx1 and TRP14 [12], which may have physiological importance.

True redox regulation of PTPN22 requires that the fully oxidized and thus inactivated enzyme can be subsequently reactivated by reduction. Since the thioredoxin system has been previously shown to reactivate phosphatases PTP1B and PTEN [11][13][16], we next tested this property. We found that H_2_0_2_-inactivated PTPN22 species could be re-activated *in vitro* using either DTT as reductant, or the thioredoxin system composed of NADPH together with TrxR1 and Trx1. Interestingly, wild-type PTPN22, but not the PTPN22^C129S^ mutant, could be directly reactivated by TrxR1 together with NADPH, without inclusion of Trx1 (Fig. 2A). Subsequent experiments showed that the effect of TrxR1-dependent reactivation was concentration dependent and that the oxidized forms of wild-type PTPN22 were clearly more amenable to direct reactivation by TrxR1 than those of PTPN22^C129S^ (Fig. 2B). Since the only difference between PTPN22 and PTPN22^C129S^ is the integrity of the non-catalytic C129 residue, we reasoned that the direct reductive reactivation by TrxR1+NADPH of wild-type PTPN22, not seen with the PTPN22^C129S^ mutant, might indicate that TrxR1 can directly reduce the disulfide involving C129 that may be formed in the wild-type PTPN22 enzyme. To assess this possibility, we subjected both forms of the enzyme to oxidative conditions, or to reduction by either DTT or TrxR1, and then analyzed the enzymes on a non-reducing SDS-PAGE. Indeed, only PTPN22 but not PTPN22^C129S^ displayed a second faster migrating form of the protein upon oxidation that disappeared upon reduction with DTT or TrxR1 (Fig. 2C). The effect was seen with several concentrations of H_2_O_2_ and DTT treatment could always revert the double band of PTPN22 (Suppl. Fig. S1). We also found that the reactivation of wild-type PTPN22, with either DTT or the thioredoxin system (TrxR1 alone, or coupled with either Trx1 or TRP14), was always more efficient than that of the mutant PTPN22^C126S^ enzyme, also in cases where prior oxidation was further potentiated by bicarbonate (Suppl. Fig. S2).

**Fig.2.**
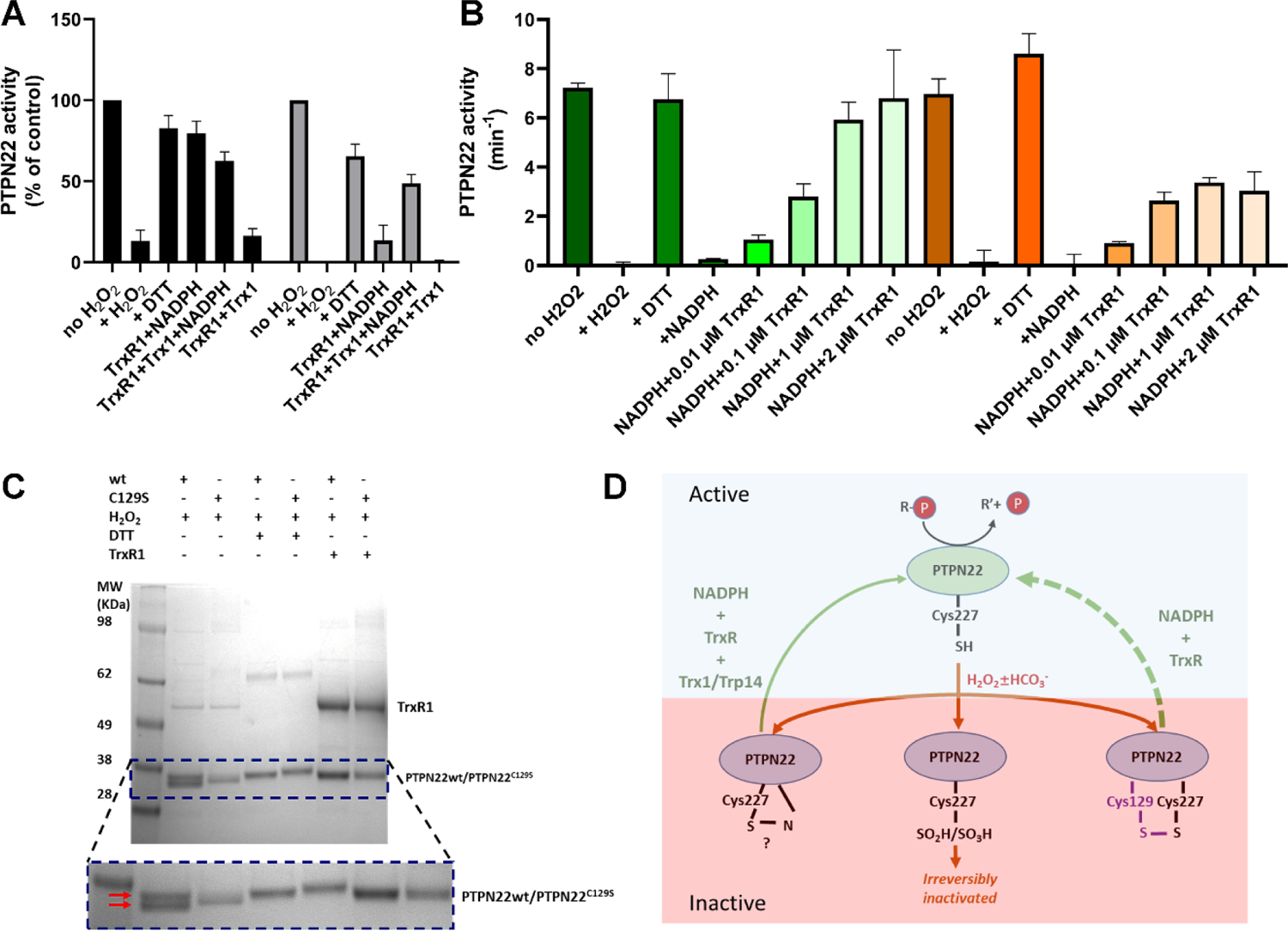
PTPN22 can be reactivated by TrxR1. **A)** Pre-reduced wild-type (black) or PTPN22^C129S^ (grey) proteins were treated with H_2_O_2_ (1mM) for 5 min whereupon residual H_2_O_2_ was removed by addition of catalase. Thereafter, the enzymes were incubated for 60 min at 37°C with either only buffer, DTT (10 mM), or with combinations of TrxR1 (2.5μM) and NADPH (300μM) and Trx1 (10μM), as indicated. Finally, the samples were analyzed for PTP activity, with activity reported as percentage of control samples treated identically but without addition of H_2_O_2_. **B)** Pre-reduced PTPN22 (green) and PTPN22^C129S^ (orange) were treated as described in A, reactivated with either 10mM DTT, or NADPH (300μM) and different concentrations of TrxR1 as indicated. After 60 min at 37 °C, samples were analyzed for PTP activity, given as turnover in min^-1^ (mol of product/mol of enzyme/min). **C)** SDS-PAGE analysis of PTPN22 or PTPN22^C129S^ treated similarly to the assay shown in A) but incubated with H_2_O_2_ (100µM), DTT (10 mM) or TrxR1 (0.5µM together with 300 µM NADPH), as indicated in the figure. The samples were subsequently analyzed on a non-reducing SDS-PAGE. Note the presence of a double band only in oxidized PTPN22, but not in oxidized PTPN22^C129S^, that disappears upon incubation with either DTT or TrxR1 (double red arrows in the enlarged portion of the gel picture). **D)** A proposed model for redox regulation of PTPN22. Only the reduced form of the enzyme is active (top, light green), which upon oxidation with H_2_O_2_ and as further facilitated by bicarbonate (HCO_3_^-^) can form several oxidized species (bottom, purple), possibly being either a sulfenylamide as suggested for PTP1B or some other species that require the complete thioredoxin system for reactivation (left), as well as irreversibly oxidized with the catalytic C227 converted to sulfinic/sulfonic acid (middle), or by forming a disulfide with C129 (right). This disulfide was here found to be amenable to reduction directly by TrxR1, but as the PTPN22^C129S^ mutant cannot form this species, it can also not be reactivated by that reduction path (dashed green arrow). Both PTPN22 and PTPN22^C129S^ can however be reactivated by the thioredoxin system from its other non-irreversibly oxidized states (solid green arrow).

These findings with recombinant human PTPN22 and PTPN22^C129S^ enzymes suggest that the mutant is more amenable to oxidation and that the thioredoxin system is less efficient in reactivating its oxidized form. The notion that TrxR1 directly can reduce the disulfide which is not formed in the mutant enzyme, but which may be formed upon oxidation of wild-type PTPN22, was interesting, as to our knowledge no PTP has earlier been shown to form a disulfide species that is a direct substrate of TrxR1. Based upon these findings, we propose a model for redox regulation of PTPN22 (Fig. 2D), which illustrates how PTPN22^C129S^ can be more easily inactivated than the wild-type enzyme. Next, we wished to assess if mutant PTPN22^C129S^ yields any phenotypic effects in mouse models of inflammation.

### Establishment of PTPN22^C129S^ mutant mice

To study redox-dependent regulation of PTPN22 *in vivo* and its possible downstream effects on inflammation we generated a mouse with the PTPN22^C129S^ mutation (schematic in Fig. 3A). This disrupts the mechanism by which the catalytic C227 can be protected through formation of a disulfide that can be reduced by TrxR1 (Fig. 2D), thus making PTPN22^C129S^ in the mice more prone to inactivation by oxidation. The mice were backcrossed to C57Bl6/N mice together with a major histocompatibility complex (*MHC*) region containing an *Aq MHC class II* allele, making it susceptible to autoimmune arthritis [15], and also with the *m1j* mutation of *Ncf1* [16], allowing interaction studies.

**Fig.3:**
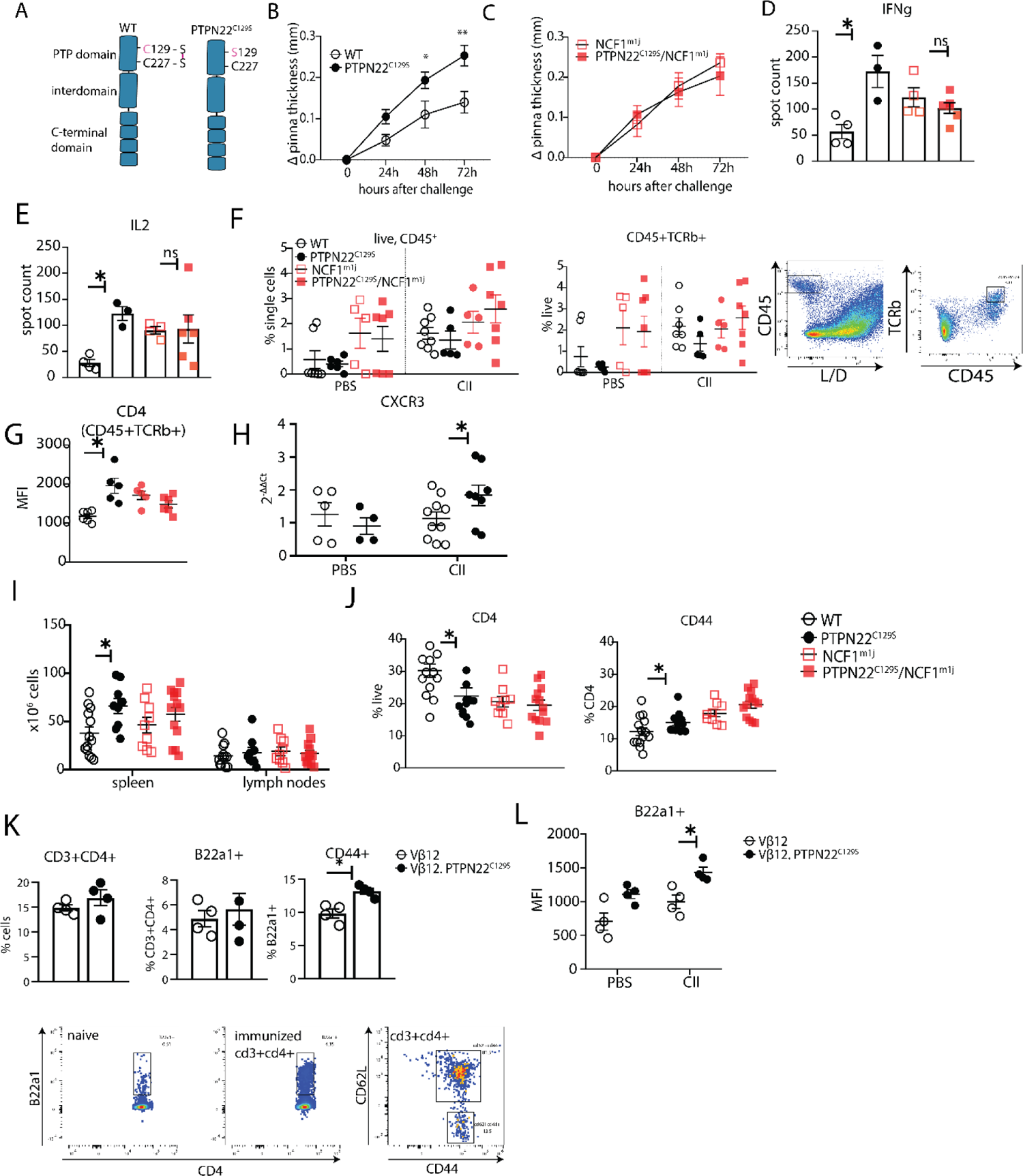
PTPN22C129S enhances Th1-mediated inflammation via NOX2-mediated ROS. **A)** Schematic representation of wild-type and PTPN22^C129S^ proteins; in the wild-type protein the catalytic cysteine 227 may form a disulfide bond with cysteine 129, whilst in the mutant, cysteine 129 is mutated into a serine, abrogating this disulfide bond. **B-L)** Mice were immunized according to DTH protocol with collagen type II and cell populations/ recall responses were analyzed 72 hours after challenge. **B-C)** Ear swelling of littermate wild-type (n=8), PTPN22^C129S^ (n=6), NCF1^mi1j^ (n=5) and PTPN22^C129S^/NCF1^m1j^ (n=7) mice at indicated time points after CII challenge. Swelling shown as mean ± SEM (2way ANOVA). Data is representative of four experiments. **D-E)** Measurement of antigen-specific interferon-γ (IFNγ)/ IL-2 T cell response against the immunodominant T cell epitope (rCII259-273) of collagen type 2 as measured by ELISpot in inguinal lymph nodes. **F)** Frequencies of indicated cell subsets in PBS-injected and CII-injected ears measured by flow cytometry; representative gating shown **G)** Mean fluorescence intensity (MFI) of CD4 within CII-injected ears measured by flow cytometry **H)** qPCR analysis of CXCR3 expression in PBS and CII-injected ears **I)** Cell count in peripheral lymphoid organs at termination of DTH protocol **J)** Frequencies of indicated cell subsets within inguinal lymph nodes measured by flow cytometry **K)** Frequencies of indicated subsets in inguinal lymph nodes of Vβ12/ Vβ12.PTPN22^C129S^ mice; flow cytometry plots show expansion of B22a1+ T cells upon immunization and representative gating for CD62L and CD44 **L)** MFI of B22a1 in PBS/CII-injected ears. Error bars represent mean ± SEM.

### PTPN22^C129S^ enhances Th1-mediated inflammation via NOX2-mediated ROS

To study the effect of PTPN22^C129S^ on cell-mediated immunity we used the delayed-type hypersensitivity (DTH) model, which is known to drive inflammation via IFNɣ producing type 1 T helper cells (Th1) [17]. Littermate wild-type and PTPN22^C129S^ mice were immunized with heterologous collagen type II (COL2), challenged on day 8 by injection with the same antigen in the ear and subsequently, swelling was assessed. We observed increased ear pinna thickness in PTPN22^C129S^ mice 24, 48 and 72 hours after challenge compared to littermate wild-type mice (Fig. 3B). In contrast, PTPN22^C129S^ did not enhance ear swelling in NCF1 mutant mice with a lack of NOX2 activity (Fig. 3C)[18]. To assess the antigen-specific T cell response we re-stimulated cells from inguinal lymph nodes with the immunodominant COL2 peptide (259-273) and observed a marked increase in IFNɣ and IL-2 producing PTPN22^C129S^ cells when comparing wild-type supporting a Th1 phenotype (Fig. 3D, 3E). No difference was observed on the NCF1 mutant background. Within the ears, flow cytometry analysis showed that the challenge with COL2 increased percentages of CD45+ leucocytes and CD45+TCRb+ cells as compared to PBS injection with the NCF1 mutants showing higher cell infiltration even in PBS-injected ears (Fig. 3F). Additionally, we observed higher CD4 expression in COL2-injected PTPN22 ears as compared to wild-type with no difference in the NCF1 mutants (Fig.3G). To further support the Th1 phenotype we performed qPCR analysis of the ears which showed increased expression of CXCR3 in inflamed PTPN22^C129S^ ears (Fig. 3H).

In the periphery, immunized PTPN22^C129S^ mice had increased cell numbers in the spleen as compared to wild-type (Fig. 3I). Within the inguinal lymph nodes we observed a reduction in CD4+ T cells which expressed higher levels of CD44, a marker for effector-memory T cells (Fig.3J). This was not seen in circulating T cells within the blood, however there was a significant reduction in FOXP3+ T cells in PTPN22^C129S^ mice 48 hours after initial immunization (Fig. S3A).

As we observed differences in T cell activation levels, we wanted to address the role of PTPN22^C129S^ in antigen-specific T cells. To do this, we used the Vβ12-transgenic mouse (Vβ12-tg) which expresses a TCRβ chain specific for the galactosylated COL2 (260-270) epitope and can be tracked using the clonotypic B22a1 antibody. Upon priming of Vβ12-tg mice with COL2 these T cells expand 50-fold, acquire an activated phenotype and play an important role in the early phase of the arthritogenic immune response [19]. Upon immunization with COL2, PTPN22^C129S^ did not affect the expansion of antigen-specific Vβ12+ T cells per se (Fig. 3K), but did change their activation status: COL2 reactive T cells in the periphery expressed higher levels of CD44 on CD4+ T cells as compared to wild-type (Fig. 3K). In the DTH model, we could also observe higher B22a1 expression among CD4+ T cells in the inflamed ears of Vβ12.PTPN22^C129S^ mice (Fig. 3L).

Together, these results indicate that the oxidation prone PTPN22^C129S^ mutant enhances T cell-mediated inflammation. Conversely, wild-type PTPN22 with a higher basal activity and more resistance to oxidation counteracts these inflammatory processes.

### PTPN22^C129S^ enhances T cell responses and development of arthritis

As PTPN22 is heavily associated with the development of autoimmunity we also sought to explore the effects of PTPN22^C129S^ on arthritis development by using the glucose-6-phosphate-isomerase peptide (GPIp)-induced arthritis model (GIA), which causes acute autoimmunity that resembles the early stages of rheumatoid arthritis, and which is regulated by NOX2-derived ROS [20, 21][21]. As shown in Fig. 4A, we observed increased arthritis severity in PTPN22^C129S^ mice from day 10 up to day 22 after onset as well as increased incidence in mutant mice. On the NCF1 deficient background we observed a trend in the opposite direction with PTPN22^C129S^ mice showing less disease and no significant difference in disease incidence (Fig. 4B).

**Fig.4:**
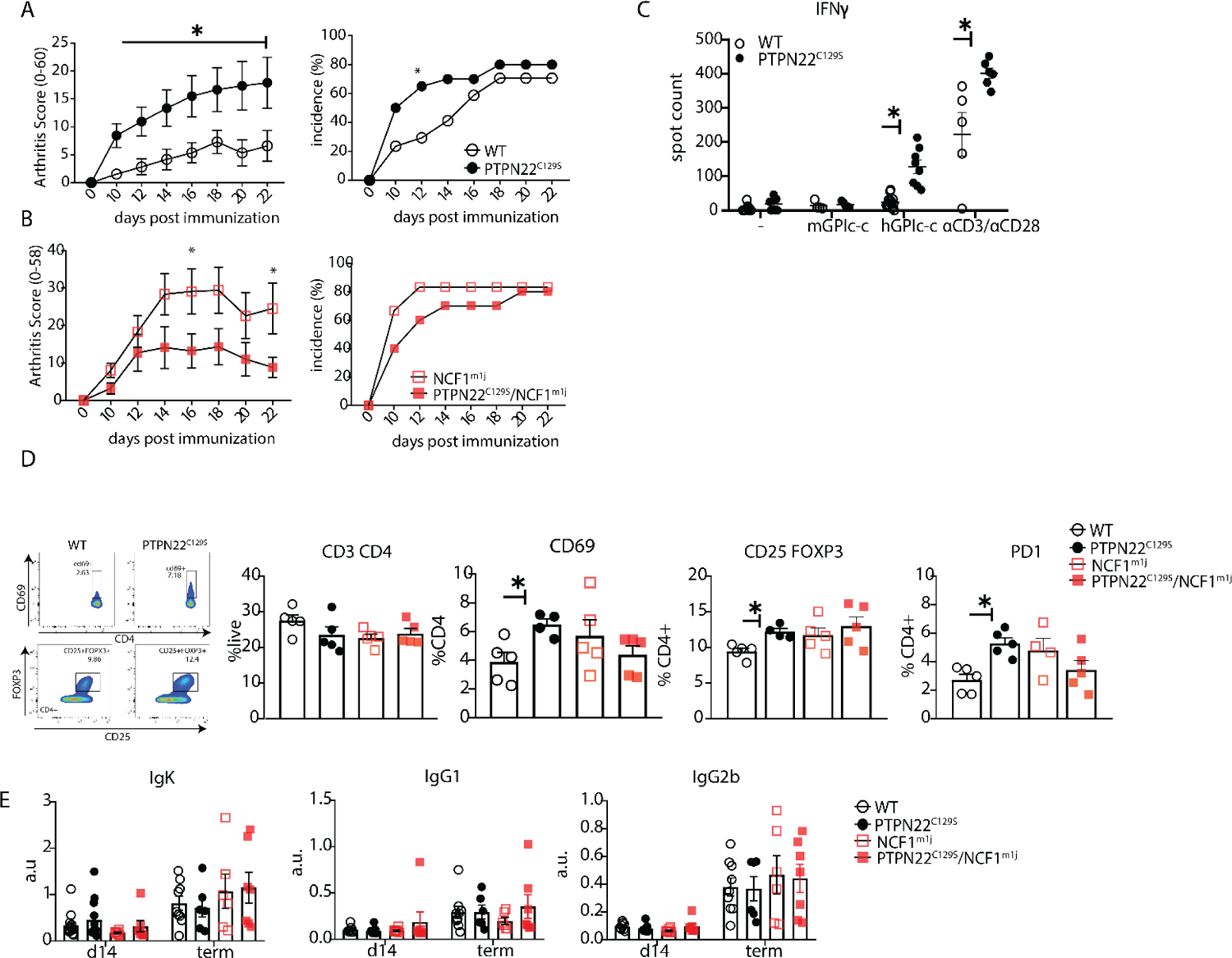
PTPN22C129S enhances arthritis development Arthritis was induced using human glucose-6-phosphate isomerase peptide (GPIp) and inflammation was monitored over 22 days. **A-B)** Clinical score (mean ± SEM) and incidence of littermate WT (n=17), PTPN22^C129S^ (n=20), NCF1^m1J^ (n=12) and PTPN22^C129S^ / NCF1^m1J^ (n=10) mice immunized with GPIp. Data from two pooled experiments (2way ANOVA). **C)** Antigen-specific interferon-ɣ (IFNɣ) T cell response against mouse/human GPIp as well as anti-CD3/anti-CD28 in lymph nodes by ELISpot **D)** Percentages of indicated cell subsets in lymph nodes at measured by flow cytometry; representative gating for CD69 and CD25+FOXP3+ cells shown. **E)** Levels of serum antibodies against hGPI day 14 post immunization and at arthritis endpoint measured by ELISA. Error bars represent mean ± SEM.

Increased disease was further supported by higher antigen-specific T cell responses in PTPN22^C129S^ mice as shown by increased numbers of IFNɣ-secreting T cells, both upon GPIp and anti-CD3/anti-CD28 stimulation (Fig. 4C). CD4 T cells in the draining lymph nodes exhibited increased activation as evidenced by higher numbers of CD4+CD69+, CD4+CD25+FOXP3+ and CD4+PD1+ T cells (Fig. 4D). No differences were observed in serum antibody levels against the hGPI peptide (Fig. 4E). In summary, PTPN22^C129S^ enhances autoimmune inflammation due to enhanced T cell reactivity.

### PTPN22^C129S^ enhances T cell signaling *in vitro* and slightly affects thymic T cell development

To understand where the observed T cell activation phenotype in PTPN22^C129S^ mice originates from, we first confirmed that PTPN22 expression was comparable between wild-type, PTPN22^C129S^, NCF1^m1J^ and PTPN22^C129S^/NCF1^m1J^ mice, both via mRNA and protein analysis (Fig. 5A-B; Fig. S3B). Next, we analyzed expansion of naïve immune cell populations. PTPN22^C129S^ was found to not affect immune cell populations in peripheral lymphoid organs in naïve mice; the percentages of B cells (B220^+^), T cells (CD3^+^), dendritic cells (CD11c^+^) and macrophages (CD11B^+^) were comparable to wild-type littermates as was MHC II expression on antigen-presenting cells (Fig. S3C).

**Fig.5:**
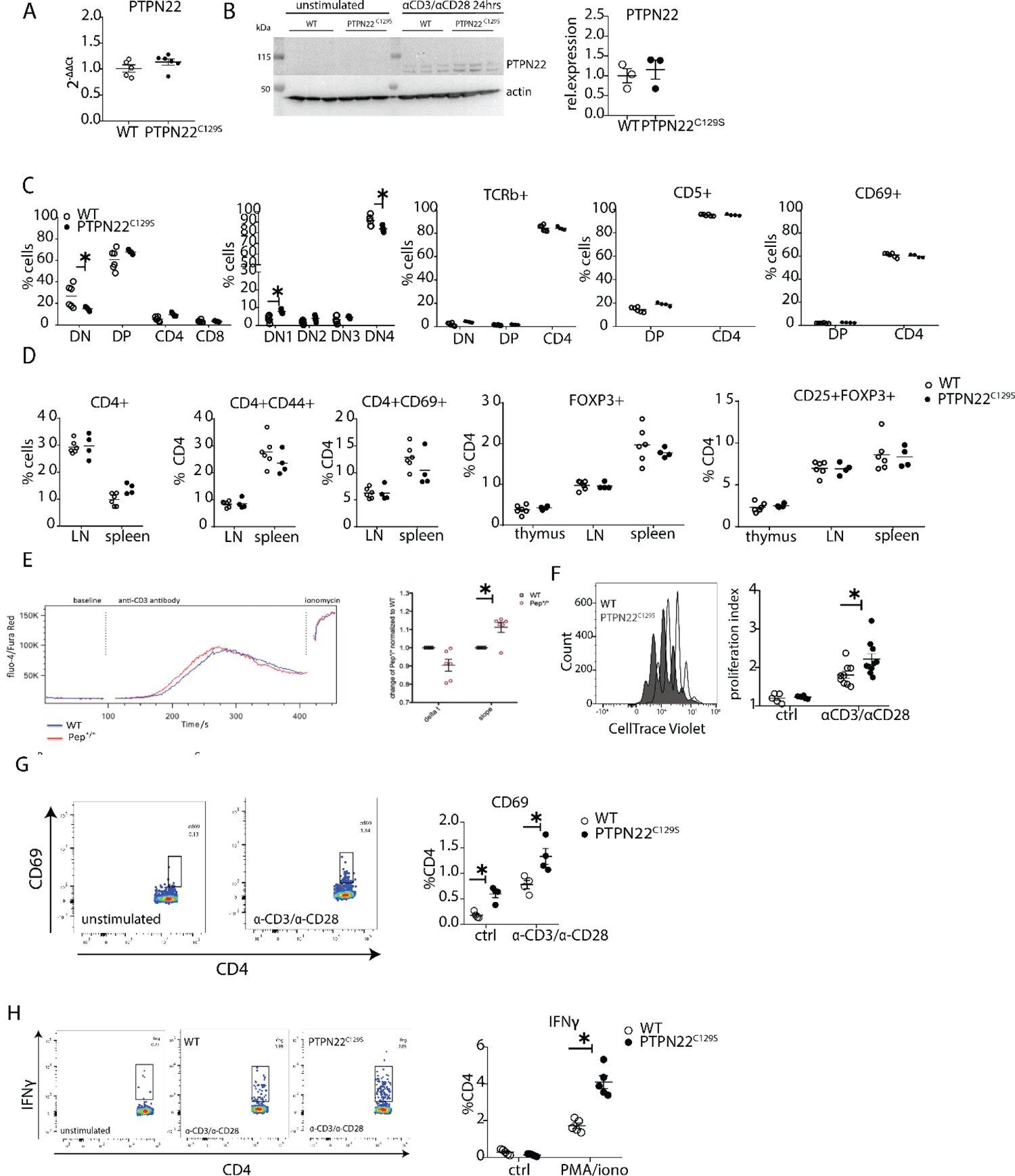
Enhanced activation and proliferation of PTPN22^C129S^ T cells. **A)** Gene expression of PTPN22 in splenocytes shown as fold change over WT. **B)** Immunoblot analysis of PTPN22 expression in sorted CD4 T cells either unstimulated or activated *in vitro* with 1ug/ml anti-CD3/CD28 for 24 hours. Each lane shows cells from a distinct mouse and relative expression to the right was calculated as by normalizing to loading control (actin) **C-D**) Analysis of T cell populations in naïve mice via flow cytometry. **C)** Analysis of double negative (DN), double positive (DP) and single positive CD4 and CD8 thymic populations in 10-week old littermate mice as well as expression of TCRb, CD5 and CD69 on various subsets **D)** Frequencies of activated (CD44+/ CD69+) and regulatory (FOXP3+) CD4+ T cells in thymus and secondary lymphoid organs. **E)** Intracellular calcium measurement in CD4 T cells at baseline, after stimulation with anti-CD3, and ionomycin to achieve maximum Ca^2+^ influx via staining with Fluo4 and FuraRed. Shown is a representative image of the change in ratio of fluo4 to FuraRed expression over time (blue=wild-type; red= PTPN22^C129S^). Quantification to the right shows slope value which describes how fast the peak of Ca^2+^ influx is reached [77] **F)** Proliferation of CD4+ T cells as assessed by CellTrace Violet dilution after 96-hour *in vitro* stimulation with 1ug/ml anti-CD3/CD28. Proliferation Index is the total number of divisions divided by the number of cells that went into division. Representative proliferation peaks on the left (clear=wild-type, black= PTPN22^C129S^) and quantification on the right (ctrl refers to unstimulated samples). **G-H)** *Ex vivo* stimulated CD4 T cells were assessed for CD69 expression/ IFNγ production after stimulation with anti-CD3/anti-CD28 or PMA/ionomycin, respectively. Representative gating shown. Error bars represent mean ± SEM.

As PTPN22 has been previously shown to affect T cell tolerance [14][15], we analyzed T cell development in the thymus. We observed a slight reduction in the percentage of CD4^-^CD8^-^ (DN) population whilst CD4^+^CD8^+^ (DP), CD4^+^, and CD8^+^ were comparable between groups. Further fractionation of the early CD4^-^CD8^-^ progenitor population showed a slight increase in DN1 (CD44^+^CD25^-^) and decrease in DN4 (CD44^-^CD25^-^) populations in PTPN22^C129S^ thymi (Fig. 5C). No changes were observed in TCRb, CD5 and CD69 expression (Fig. 5C). Percentages of CD4^+^ and CD8^+^ T cells as well as expression of CD44, and CD69 activation markers on CD4+ T cells were not affected (Fig.5D). Furthermore, percentages of CD4+FOXP3+ and CD4+FOXP3+CD25+ cells were comparable between groups in thymus and peripheral lymphoid organs. (Fig. 5D).

As we consistently observed a differential T cell phenotype in mice with mutated PTPN22^C129S^ *in vivo*, we studied T cell function *in vitro*. To assess signaling, we measured Ca^2+^-flux, one of the earliest signaling events upon TCR engagement and observed slighly increased intracellular Ca^2+^ in PTPN22^C129S^ CD4 T cells upon anti-CD3 stimulation (Fig. 5E). Proliferation of CD4 T cells, as assessed by CellTrace dilution, was also increased upon anti-CD3/antiCD28 stimulation (Fig. 5F). We observed increased activation of *ex vivo*-stimulated CD4 T cells as measured by the early T cell activation marker CD69 as well as markedly different IFNɣ production resembling the *in vivo* results (Fig.5G-H).

### Enhanced phosphorylation of PTPN22 targets and enhanced PKC expression in PTPN22^C129S^ cells

To assess whether the differences in T cell reactivity could be correlated with altered PTPN22 catalytic activity we analyzed the phosphorylation of its targets Fyn, LCK and Zap70. TCR-stimulation of wild-type and mutant cells revealed increases in phosphorylation of tyrosines Y420 and Y394 of the Src family kinases Fyn and LCK, respectively as well as of Y493 of Zap70, thus agreeing well with lower intracellular phosphatase activity of the PTPN22^C129S^ mutant protein (Fig. 6A-B) and confirming findings with recombinant PTPN22 proteins.

**Fig.6:**
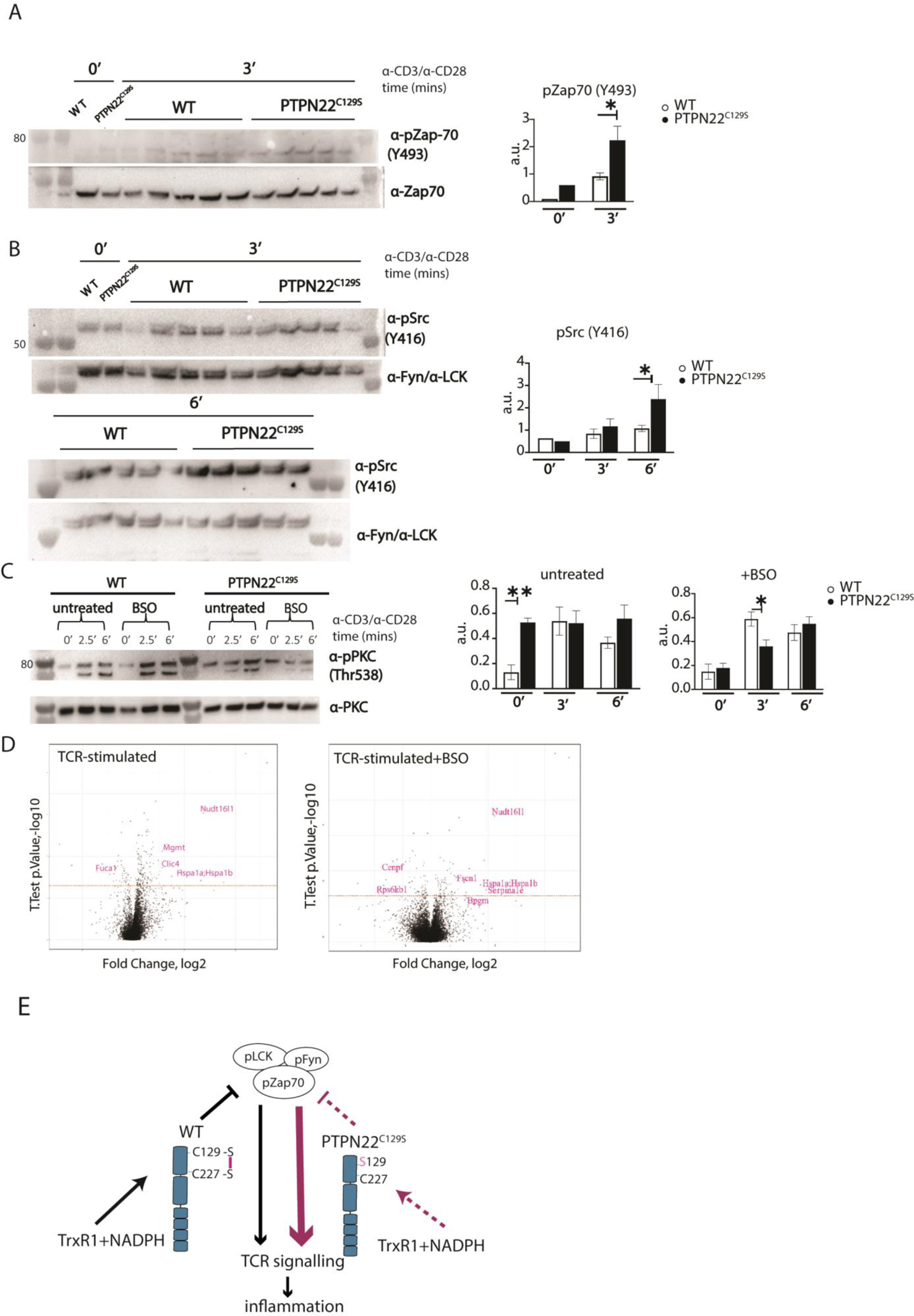
Enhanced phosphorylation of PTPN22^C129S^ targets and increased PKCθ expression. **A-B)** Lymph node cells were stimulated for indicated time points with anti-CD3 plus anti-CD28 and phosphorylation was detected via immunoblotting of total cell lysates with antibodies against activated Zap-70 (A) and Src kinases (B). Antibody against p-Src (Y416) binds to Lck Y394 and Fyn Y417 with the top band representing pFyn and the bottom band pLCK, see [78]. Each lane reflects distinct wild-type/ PTPN22^C129S^ mice and unlabeled lanes show molecular weight markers. Quantification to the right was calculated as phosphorylation signal/ total signal normalized to loading control. **C)** Lymph node cells were cultured 24hours with/ without BSO, a glutathione depletion agent, before *in vitro* activation with anti-CD3 plus anti-CD28 for indicated time points. Total lysate was analyzed for PKCθ expression via immunoblotting. Representative image shown. Quantification to the right was calculated as phosphorylation signal/ total signal normalized to loading control (WT=6, PTPN22^C129S^=8). **D)** Volcano plots comparing the proteomic profile of sorted CD4+ T cells from wild-type and PTPN22^C129S^ mice which were stimulated with anti-CD3 plus anti-CD28 for 5 hours either with or without prior treatment (24h) with BSO. Four mice per group were used for proteomic analysis and p-values were calculated by Welch’s t-test. Proteins that were significantly up/downregulated in PTPN22^C129S^ mutants are highlighted in red. **E)** A proposed model for redox regulation of PTPN22 *in vivo*: upon oxidative pressure within the cell C227 of PTPN22 forms a disulfide bond with the back-door cysteine C129 as a means of protection from irreversible oxidation and inactivation. When the C227-C129 bond cannot be formed, as is the case in PTPN22^C129S^ mutant, PTPN22 is less sensitive to reactivation by the thioredoxin system which leads to increased TCR signaling and inflammation. Error bars represent mean ± SEM.

Next, we wanted to assess if downstream T cell-signaling mediators were affected by PTPN22^C129S^. TCR-signaling induces activation of protein kinase C (PKC)-θ, an essential player in peripheral T cell activation. As compared to wild-type, PTPN22^C129S^ lymph node cells showed constitutive phosphorylation of Y538 on PKC-θ, independent of TCR stimulation whereas in wild-type cells phosphorylation increased with TCR stimulation, hinting at higher basal activity of PKC-θ in the mutant (Fig. 6C). To assess how changes in the redox status affect T cell signaling we used buthionine sulphoximine (BSO), an inhibitor of ɣ-glutamyl-cysteine synthetase, which is essential for glutathione synthesis. Treatment with BSO increases the oxidative burden in cells without directly affecting the thioredoxin system. Here, it initially reverted the phospho-PKC-θ levels in the mutant to be comparable with wild-type, followed by a slower TCR-activation-dependent phosphorylation (Fig. 6C). Next, we investigated how differential signaling was affecting the proteome signature using sorted CD4+ T cells from wild-type and PTPN22^C129S^ mice. Mass spectrometric proteomics analysis of activated cells revealed differential expression of multiple genes under both untreated and BSO-treated conditions: Up-regulation of *Nudt16l1* [18, 19], *Clic4* [26], *Mgmt* [27], *Hsp* [28], *Serpina1e* [29] and *Fscn1* [30] as well as down-regulation of *Fuca1* [31] and *Rps6kb1* [32] have all been shown to be associated with increased PKC signaling (Fig. 6D).

Thus, PTPN22^C129S^ that is prone to inactivation by oxidation with decreased catalytic activity results in enhanced T cell signaling, which has broad signaling effects that can yield aggravated inflammatory disease.

## Discussion

In this study we investigated whether redox regulation of PTPN22 has an impact on inflammation and autoimmune arthritis. We found that the C129S mutation in *Ptpn22* leads to reduced catalytic activity and increased susceptibility to oxidative inactivation *in vitro*, causing concomitant upregulation of T-cell activation and T-cell-mediated autoimmunity in a NOX2-dependent manner.

Redox regulation describes the modulation of signaling pathways by oxidation and reduction, the effects of which must always be considered in relation to enzymatically driven oxidative and reductive pathways, and the signaling properties of redox-modified cellular targets [33][34]. Significant ROS producers include the membrane-bound NADPH oxidase (NOX) complexes as well as the mitochondrial respiratory chain, modulating responses such as cell growth, death and immune function under both physiological and pathological conditions [35]38]. The interplay between redox signaling and the immune response is complex and multi-faceted. The protective effects of ROS against inflammatory diseases *in vivo* is best evidenced in the *Ncf1^m1J^* mouse where diminished ROS leads to exacerbated inflammation in arthritis and lupus models [18][37]. In T cells, ROS, most likely derived from NOX2 expressing antigen presenting cells have been shown to regulate NFAT activation, IL2 production and plasticity by both influencing the redox status of kinases involved as well as through metabolic reprogramming [8][38][39]. The list of intracellular signaling molecules regulated by ROS is long: transcription factors such as Nrf2/Keap1, NF-kB, HIF-1a, AP-1 as well as key TCR downstream mediators such as MAPK/ERK, LCK, Zap70, LAT, PKC-θ and PLCγ1 are all redox-sensitive [40]. The impact of the thioredoxin- and glutathione-dependent reductive systems on these signaling pathways is also of crucial importance [34][41].

Reversible cysteine modifications have emerged as a major redox regulatory mechanism important for ERK1/2 phosphorylation, calcium flux, cell growth, and proliferation of naïve CD4+ and CD8+ T cells [42]. Proteins containing cysteine residues, such as protein tyrosine phosphatases (PTPs), are particularly susceptible to redox effects in part due to the low pKa value of their active site. Oxidation leads to inhibition of PTP activity and several PTPs such as Src homology region 2 domain-containing phosphatase-1 and 2 (SHP1/2) and tyrosine-protein phosphatase non-receptor type 1 (PTP1B) are well studied in terms of their redox regulation [9][43][44][45]. Again, the balance between oxidative and reductive pathways is crucial for overall PTP activities and in this context the thioredoxin system is important for its reductive PTP-activating capacity [13][46][47].

PTPN22 is a protein tyrosine phosphatase expressed in all hematopoietic cells and a known negative regulator of T cell signaling. The R620W variant of PTPN22 has been associated with an increased risk of several autoimmune diseases, amongst them diabetes, rheumatoid arthritis and systemic lupus erythematosus. It has been shown to interfere with the binding of PTPN22 to the C-terminal SRC kinase (CSK) affecting downstream signaling. However, whether R620W is a gain-of-function or loss-of-function variant remains controversial [24][25][48]. A possibility for redox regulation of PTPN22 was highlighted by Tsai *et al.* who discovered an atypical disulfide bond formation between the catalytic cysteine C227 and a “back-door” cysteine C129, as visualized in a crystal structure of the catalytic domain of PTPN22 [10]. As physiologically relevant oxidative regulation of PTPN22 has thus far not been studied, we decided to mutate C129 into a serine, thereby abrogating the capacity of C129 to contribute to redox regulation of PTPN22, both *in vitro* and *in vivo*.

In our *in vitro* experiments, PTPN22^C129S^ showed a lower basal catalytic activity, higher sensitivity to inhibition by oxidation, and a lower capacity for reactivation by the thioredoxin system, compared to the wildtype enzyme. The reason behind the initial drop in activity remains to be studied; the single cysteine-to-serine change may affect the PTPN22 protein conformation or dynamics and crystallography studies would be needed to address this. It should also be noted that the pure PTPN22^C129S^ migrated slightly slower than wild-type PTPN22, despite the only difference between the two proteins being the single C129 residue. This also suggests an overall effect on protein characteristics by the C129S mutation, which should be studied further. Based on our findings, we propose a model for the redox regulation of PTPN22, with the disulfide bond that can be formed between C227 and C129 protecting the catalytic C227 residue from overoxidation to sulphinic or sulphonic acid. It is also possible that an alternative sulphenylamide motif can be made between the thiol group of the C227 with the peptide bond amine, which can also be reduced by the thioredoxin system, similar to that seen with PTP1B [13][14] even if that motif was not seen in the crystal structure of PTPN22 [10]. This model explains how PTPN22^C129S^ can still show activity and a certain protection against oxidation and reactivation by the thioredoxin system, but simultaneously be more susceptible to oxidation because the protective disulfide cannot be formed.

In T cells, the PTPN22^C129S^ mutation led to enhanced T cell activation and proliferation. This was not due to altered PTPN22 expression levels, but rather a consequence of downstream signaling. The PTPN22 targets LCK, Fyn and Zap70 showed increased phosphorylation, arguing for lower catalytic activity of PTPN22^C129S^ *in vivo* and confirming our *in vitro* findings. In TCR signaling, phosphorylation of PKC-θ is a late-stage event that leads to the activation of transcription factors Nf-Kb, NFAT and AP-1 controlling T cell functions down the line [49]. We observed baseline upregulation of p-PKC-θ in PTPN22^C129S^ T cells which did not respond to further TCR stimulation. PKC phosphorylation is regulated by GLK (germinal center kinase-like kinase) and LCK [49]–[51]. An increase in LCK phosphorylation as previously observed might therefore be causing increased p-PKC-θ. In an oxidative microenvironment created by BSO treatment, the initial difference in unstimulated cells disappeared and PTPN22^C129S^ instead showed slower phosphorylation kinetics compared to wild-type. This might be mediated by direct effects of ROS on PKC as it contains several redox-sensitive residues that can lead to PKC inactivation [52]. Proteomics data further corroborated the enhanced activation state. Interestingly, in the *Ptpn22* mutant, we observed down-regulation of CENPF, a cell cycle protein that was shown to be upregulated upon TrxR inhibition [53].

Enhanced T cell activation led to increased inflammation *in vivo* and we observed a pronounced skewing towards Th1 responses. A link between PTPN22 and Th1 responses has previously been established; PTPN22^-/-^ and PTPN22^R619W^ mice accumulate Th1 effector cells in their lymphoid organs with age or upon immune challenge. Th1 expansion in these mice was found to be dependent on LFA-ICAM1 interactions [54]. A collagen-induced arthritis (CIA) model produced no visible differences in inflammation in PTPN22^C129S^ mice, in line with previous data showing that even a complete lack of PTPN22 does not result in significant differences in inflammation (Supp. Fig.3D) [54]. An explanation could be that the major pathogenic factor in the CIA model are autoreactive B cells producing pathogenic antibodies, rather than autoreactive T cells [55].

In both T cell mediated inflammatory models (DTH and GPI) as well as on the COL2-T cell transgenic background, PTPN22^C129S^ mice displayed higher T cell activity as well as more regulatory T cell markers such as FOXP3 and PD1 (shown for GPI model). A rise in Treg numbers is a measure to combat inflammation, but where they originate from is a matter of discussion. They can originate from the thymus and migrate to the tissue in response to inflammation or result from conversion of CD4+CD25-naïve T cells in the periphery [56]. PTPN22 plays an important role in determining TCR signal strength during central tolerance; PTPN22-deficient mice show increased positive selection in the thymus as well as increased Treg numbers both in the thymus and periphery leading to protection from the experimental autoimmune encephalomyelitis (EAE) model of autoimmunity [18][44][45]. In our study, Treg numbers were comparable between wild-type and PTPN22^C129S^ thymi, suggesting peripheral induction of Tregs upon inflammation in PTPN22^C129S^ mice. This may be a consequence of increased IFNɣ production by PTPN22^C129S^ T cells as studies have also associated FOXP3 induction to Th1 responses; this mechanism links IFNɣ and IL27 with amplifying TGF-b-induced FOXP3 expression via STAT1 [61][62]. Previous studies have also shown the essential role of PKC-θ in NFAT-dependent FOXP3 expression. PKC-θ^−/−^ mice have significantly reduced CD4+ FOXP3+ Tregs in the thymus, spleen and lymph nodes [51]. Accordingly, the rise in Tregs in PTPN22^C129S^ mice may be a consequence of the enhanced baseline PKC-θ signaling.

In summary, our results show how the activity of PTPN22, a gene where a risk variant is associated with autoimmune diseases, can be regulated by ROS through its non-catalytic C129 residue, which is likely to have a major impact on its function in addition to that of merely altered basal turnover (Fig.6D). It has been notoriously difficult to target PTPs *in vivo* with drug therapies due to the nature of their active sites [61], but recent advances in targeting the redox status of T lymphocytes might represent a novel strategy to treat T-cell-driven diseases. Antibody-trapping of the oxidized form of PTP1B, a highly touted drug target for diabetes and obesity, has been shown to increase insulin signaling *in vitro* [6][62][63]. It is therefore of importance to study the redox regulatory effects on the T cell machinery and its effects on immune processes. As such, our study contributes to the understanding of redox regulation of PTPN22 and possibly represents a new avenue for targeting PTPN22.

## Materials and Methods

### Recombinant expression of PTPN22

The open reading frame of human PTPN22 catalytic domain (1-303 residues), codon optimized for recombinant expression in *E. coli*, was synthesized by Integrated DNA Technologies, Inc. The ORF was subcloned into an in-house developed pD441b plasmid (pD441b-HsPTPN22cd) that generates a fusion protein of His6-sfGFP-SUMO-HsPTPN22cd, where His6 is an N-terminal His-tag for IMAC purification, sfGPF (superfolder GFP, [64]) included for enhanced protein folding, solubility and direct visibility, and SUMO (small ubiquitin-related modifier) being a 110 residue sequence recognized by SUMO protease ULP1 that hydrolyzes the peptide bond at the C-terminus of the SUMO domain, resulting in release of the target HsPTPN22cd from its N-terminal fusion partner. The C129S mutation of human PTPN22 was introduced into the wild-type PTPN22-encoding plasmid pD441b-HsPTPN22cd by PCR using primer pairs HsPTPN22-C129S-fwd (GATCGGTCTCATGGCGAGCATGGAGTACGAGATGGG) and HsPTPN22-C129S-rev (GATCGGTCTCCGCCATAACAATGATCAGTAC), and the resulting plasmid was named pD441b-HsPTPN22cd-C129S. Both plasmids were respectively transformed into E. coli BL21(DE3) strain for propagation and protein expression. The full sequences of the open reading frames of the constructs for the wild-type PTPN22cd and its C129S mutant were verified by sequencing (Eurofins Biotech). Briefly, for each protein expression and purification, a 40 mL overnight culture was inoculated into 2L terrific broth (TB) medium containing 50 µg/mL kanamycin in a 5L-bottle placed on a shaking incubator at 37 °C. Four hours after the inoculation, the incubator temperature was lowered to 25 °C and 0.5 mM IPTG was added to induce protein expression overnight. The bacteria were harvested by centrifugation, suspended in IMAC binding buffer (50 mM Tris-HCl, 100 mM NaCl, 10 mM imidazole, pH 7.5) and lysed by sonication. The soluble fraction was recovered by centrifugation and purified by applying a HisPrep FF 16/10 column used with an ÄKTA explorer FPLC system (Cytiva Life Sciences). The eluted and purified fusion protein was subsequently treated with an in-house expressed and purified His-tagged ULP1 (1% w/v) and the cleavage solution was again re-applied onto the HisPrep FF 16/10 column for separation of non-tagged target protein from its N-terminal His-tagged fusion partner as well as from the His-tagged ULP1. The target protein was then concentrated, buffer exchanged and stored in the freezer. The purity of the product was greater than 95% as assessed by SDS-PAGE.

### PTP activity assay

PTP activity of recombinant wild-type PTPN22 and PTPN22^C129S^ (1400nM) was measured spectrophotometrically using as substrate 15mM chromogenic *p*-nitrophenyl phosphate (pNPP) (P4744-1G, Sigma-Aldrich), as described previously [65]. The absorbance increase rates were measured at 410 nm at 22 °C using an Infinite M200 Pro plate reader (Tecan). Reduced PTPN22 (1400nM, 49.9μg/ml) was preincubated in 20 mM HEPES, 0.1mM diethylenetriaminepentaacetic acid and 100 mM NaCl buffer (pH 7.4) containing 0.05% BSA to prevent PTPN22 from time-dependent inactivation [11] together with 1 mM sodium azide. Sodium azide was used to inhibit any trace amounts of catalase. At the indicated concentrations human Trx1, TrxR1 and NADPH (N7505-100MG, Sigma-Aldrich) were added (with Trx1 and TrxR1 expressed and purified as described in [66]. Variations in activity were observed between different batches of PTP purifications, and activities were thus always compared with the controls within each experiment.

### PTP treatment with H_2_O_2_ and bicarbonate

Pre-reduced and desalted PTPN22 was exposed to H_2_O_2_ and different components of the Trx system at the indicated time points followed by the addition of pNPP and measurement of activity. The activity after each H_2_O_2_ treatment was related to the activity of untreated PTPN22 incubated for the same time. For analysis of the effects of bicarbonate, similarly pre-reduced PTPN22 (1400nM) was pre-incubated in 20mM HEPES, 100 mM NaCl buffer, pH 7.4, containing 0.1mM EDTA, 0.05% BSA. For treatment with oxidant, bicarbonate and H_2_O2 were premixed in the same buffer before addition to PTPN22 for treatment. Each PTPN22 in reaction mixture was exposed for the indicated times to H_2_O_2_ with or without bicarbonate (pH 7.4) and subsequent to this treatment, measurement of PTP activity was performed.

### PTP reactivation assay

PTPs from stock storage solutions were exchanged into reactivation buffer as described previously [67] containing 0.5% (v/v) Tween 80. Subsequently the PTPs were inactivated by treatment with 1 mM H_2_O_2_ for 5 min at 25°C. Following this inactivation incubation, 20 μg/ml catalase was added to quench excess H_2_O_2_ whereafter the reactivation experiment was performed by adding the components of the Trx system as indicated.

### Animals

#### Establishment of mouse strains

##### Vector construction

A genomic fragment encompassing exon 2 to 10 of the *Ptpn22* gene was isolated from a BAC clone of C57BL/6 origin (n° RP23-189D14, Deutsches Ressourcenzentrum für Genomforschung). The TGT codon found in exon 5 of the *Ptpn22* gene and coding for the cysteine residue present at position 129 of PTPN22 was converted into a TCC codon coding for a serine. A *lox*P-tACE-CRE-PGK-gb2-*neo*r-*lox*P cassette (NEO;[68]) was introduced in the intron separating exons 5 and 6 of the *Ptpn22* gene, and a cassette coding for the diphtheria toxin fragment A abutted to the 5’ end of the targeting vector.

#### Isolation of recombinant ES clones

After electroporation of Bruce 4 C57BL/6 ES cells [69] and selection in G418, colonies were screened for homologous recombination by Southern blot. A probe specific for the NEO cassette was also used to ensure that adventitious non-homologous recombination events had not occurred in the selected ES clones.

#### Production of Mutant Mice

Mice were handled in accordance with national and European laws for laboratory animal welfare and experimentation (EEC Council Directive 2010/63/EU, September 2010), and protocols approved by the Marseille Ethical Committee for Animal Experimentation. Mutant *Ptpn22*^C129S^ ES cells were injected into FVB blastocysts. Germline transmission led to the self-excision of the NEO selection cassette in male germinal cells. Presence of the intended mutation was checked in homozygous *Ptpn22*^C129S^ mice by DNA sequencing. Screening of mice for the presence of the *Ptpn22*^C129S^ mutation was performed by PCR using the following oligonucleotides: 5’-CATGCAGGACTGTCCTCTCT-3’ and 5’-GTATTCTTGTCTCCCTTCCT-3’. This pair of primers amplifies a 283 bp band in the case of the wild-type allele and a 368 bp band in the case of the *Ptpn22*^C129S^ allele. Mice harboring the *Ptpn22*^C129S^ mutation have received the international strain designation C57BL/6-*Ptpn22*^tm1Ciphe^, and founders were sent to Karolinska for further experiments.

The mice were backcrossed to C57Bl6/N.H2^q^ (B6Q) mice. This genetic background was used in all experiments. In some experiments we introgressed a TCR VDJ beta knock in as well as a TCR alpha locus originally derived from the DBA/1 mouse [70]. For other experiments, the Ncf1^m1j^ mutation was inserted, leading to a functional impairment of the NOX2 mediated ROS response [71]. Mice were kept under specific pathogen free (SPF) conditions in the animal house of the Section for Medical Inflammation Research, Karolinska Institute in Stockholm. Animals were housed in individually ventilated cages containing wood shavings in a climate-controlled environment with a 14 h light-dark cycle, fed with standard chow and water *ad libitum*. All mice were healthy and basic physiological parameters were not affected. All the experiments were performed with age-, and sex-matched mice and in a blinded fashion. Experimental procedures were approved by the ethical committees in Stockholm, Sweden. Ethical permit numbers: 12923/18 and N134/13 (genotyping and serotyping), N35/16 (DTH, GPI, CIA).

### Delayed Type Hypersensitivity (DTH)

12-week old mice were sensitized by intradermal injection of 100ug rat collagen type II (COL2) in 100ul of a 1:1 emulsion with Complete Freund’s adjuvant (BD, Difco, MI, USA) and 10mM acetic acid at the base of the tail. Day 8 after sensitization the right ear was injected intradermally with 10ul COL2 in PBS (1mg/ml) whilst the control left ear was injected with 10ul acetic acid in PBS. Ear swelling response was measured 0h, 24h, 48 and 72 hours after challenge using a caliper. Change in ear thickness was calculated by subtracting the swelling of the PBS-injected ear from the swelling of the COL2-injected ear normalized to day 0 thickness. Ear tissue was processed according to [72].

### GPI-induced arthritis (GPI)

Arthritis was induced by the hGPIc-c peptide (NH2-IWYINCFGCETHAML-OH; Biomatik) emulsified with an equal volume of complete Freund’s adjuvant. Each mouse was intradermally injected with 100ul emulsion (10ug/mouse) at the base of the tail and arthritis development was monitored using a macroscopic scoring system where visibly inflamed ankles or wrists received 5 points each and inflamed toes/fingers were given 1 point.

### Collagen-induced arthritis (CIA)

Arthritis was induced with 100 µg of heterologous rat COL2 in 100 µl of a 1:1 emulsion with CFA and 10mM acetic acid injected intradermally at the base of the tail. Mice were challenged on day 28 with 50 µg of COL2 in 50 µl of IFA (BD, Difco) emulsion. Arthritis development was monitored using a macroscopic scoring system as described above.

### Cell culture

10^6^ splenocytes or lymph node cells were cultured in 200 µl of complete DMEM per well in U-shaped bottom 96-well plates (Thermo) at 37°C and 5% CO2. Complete DMEM consisted of: DMEM+Glutamax (Gibco); 5% FBS (Gibco); 10 µM HEPES (Sigma); 50 µgml^−1^ streptomycin sulfate (Sigma); 60 µgml^−1^ penicillin C (Sigma); 50 µM beta-mercaptoethanol (Gibco). FBS was heat-inactivated for 30 min at 56°C. C. Following stimuli were used: hGPIc-c (10µM; Biomatik), mGPIc-c (10uM, Biomatik), galCOL2259–273 (10µgml^−1^, in-house), concanavalin A (ConA, 1ugml^−1^), anti-mouse CD3 (1µgml^−1^, 145-2C11, BD); anti-mouse CD28 (1µgml^−1^, 37.51, BD), Phorbol 12-myristate 13-acetate (20ngml^−1^), ionomycin (1µgml^−1^)

### ELISA

Flat 96-well plates (Maxisorp, Nunc) were coated overnight at 4°C with 10ug/ml rCOL2/GPIp-BSA in PBS. Plates were washed in PBS+0.05% Tween and blocked with 1%BSA in PBS for 2 hours at 37C and washed again. Mouse serum was added in dilutions varying from 1:100 to 1:10000. Plates were incubated for 2 h at 37C, washed and incubated with secondary antibodies (Southern Biotech: IgK (1170-05), IgG2b (1091-05)) for 1hour at 37C. After washing, 50ul ABTS substrate (Roche) was added and signal was read at 405nm (Synergy 2, BioTek).

### T cell proliferation

Proliferation of CD4+T cells purified by negative sorting (Thermo, 11416D) was assessed using the CellTrace Proliferation Kit (Thermo, C34554) according to manufacturer’s instructions.

### ELISPOT

EMD Millipore MultiScreen 96-Well Assay Plates were coated overnight at 4°C with the capture antibody in PBS. Coating solution was decanted and 10^6^ splenocytes or 5 × 10^5^ lymph node cells were incubated 24hrs at 37C together with various stimuli. After culture, plates were washed (0.01% PBS-Tween) and biotinylated detection antibodies were added in PBS. Plates were washed and Extravidin Alkaline phosphatase (Sigma) was added at a 1:2500 dilution in PBS (30 min, RT). Plates were washed before adding Sigmafast BCIP/NBT (Sigma) substrate solution. Once spots became visible, plates were washed in water and spots were counted using a CTL ImmunoSpot Analyzer. <u>Antibodies (Ab)</u>: IL-2 (capture Ab 5µgml^−1^ JES6-IA12; detection Ab 2µgml^−1^ biotinylated-JES6-5H4, in-house produced); IL-17A (capture Ab 5 µgml^−1^ TC11-18H10.1; detection Ab 2,5µgml^−1^ TC11-8H4, Biolegend); IFNɣ (capture Ab 5 µgml^−1^ AN18; detection Ab 2,5 µgml^-1^ biotinylated R46A2, in-house produced).

### qPCR

RNA was extracted from 1-2×10^6^ cells using Qiagen RNeasy columns according to the manufacturer’s instructions. RNA concentration was determined using a NanoDrop 2000 (Thermo Scientific) and sample concentrations were normalized before proceeding with reverse transcription. cDNA synthesis was carried out using the iScript cDNA synthesis kit (BioRad) according to manufacturer’s instructions. The qPCR reaction was carried out using the iQSYBR Green Mix (BioRad) in white 96-well plates using a CFX96 real-time PCR detection system (BioRad). *Actb* was used as an endogenous control. Data were analyzed according to the ΔΔCt method [73]. Primer sequences are listed in supplemental table 1.

### Flow cytometry

Samples were stained with the indicated antibodies in 50µl of PBS diluted 1:200 at 4°C for 20 min in the dark. Cells were washed once, fixed and permeabilized for intracellular staining using BD Cytofix/Cytoperm™ (BD) according to manufacturer’s instructions. Cell were stained intracellularly with 50 µl of permeabilization buffer (BD), using the antibodies at a 1:100 final dilution, for 20 min at RT. FoxP3 staining required nuclear permeabilization and was carried out using Bioscience™ Foxp3 / Transcription Factor Staining Buffer. For intracellular cytokine staining, cells were stimulated *in vitro* with phorbol 12-myristate 13-acetate (PMA) 10ngml^-1^, ionomycin 1µgml^-1^, and BFA 10µgml^-1^ for 4-6 h at 37°C prior to fixation, permeabilization and staining. Antibodies are listed in supplemental table 2.

### Ca2+ flux

Lymph node and spleen cells were stained in pre-warmed PBS+1%FCS with Fluo4-AM (2uM, Thermo) and FuraRed AM (4uM, Thermo) at 37C. Cells were washed in cold PBS+1%FCS before staining for extracellular markers for 20 mins. Baseline Ca2+ flux was recorded for 100s before 50ul anti-CD3 (10ug/ml; BD) stimulation was added; after 5 minutes, maximum flux was measured using ionomycin (1ug/ml; BD). Relative calcium concentration was plotted as ratio of Fluo3 to FuroRed emission using FlowJo.

### Protein isolation and SDS-PAGE

Total protein was isolated from 2 × 10^6^ cells in 60µl lysis buffer (M-PER, Thermo) with freshly added protease inhibitors (Halt cocktail 100x, Thermo). Lysates were centrifuged for 10 min at top speed and protein concentrations of supernatants were measured using Pierce BCA Protein Assay Kit (Thermo, 23225). SDS-PAGE (4-12% NuPAGE Bis-Tris gel; Thermo Scientific) was run according to manufacturer’s instructions (45 min, 200 V, MOPS buffer).

### Western blot

Proteins were blotted onto a PVDF membrane (Millipore) for 1,5 h at 35 V in NuPAGE transfer buffer (Thermo Fisher). Membranes were blocked for 1 h at RT in blocking solution (0.05% PBS-Tween, 5% BSA). Incubation with primary antibodies (listed below) was performed overnight at 4°C in blocking solution. After incubation, membranes were washed in PBS-T and incubated with Affinipure peroxidase-coupled goat anti-rabbit IgG(H+L) (final conc. 40 ngml^−1^, Jackson laboratories) for 1 h at RT. Membranes were washed and coated with ECL substrate solution (GE Healthcare) before imaging on a ChemiDoc XRS+ (BioRad). <u>Antibodies</u> Cell Signaling Technology: p-PKC (9377S), PKC (1364S), Zap70 (2705S), PTPN22 (14693S), p-Src (2101S), LCK (2752S), Fyn (4023S), H2B (12364s) Abcam: p-Zap70 (ab194800), actin (ab8227), vinculin (ab129002)

### LC-MS sample preparation

For proteomics analysis, cells were collected after treatment, washed twice with PBS, and then lysed using 8M urea, 1% SDS, and 50mM Tris at pH 8.5 with protease inhibitors (Sigma; Cat#05892791001). The cell lysates were subjected to 1 min sonication on ice using Branson probe sonicator and 3 s on/off pulses with a 30% amplitude. Protein concentration was then measured for each sample using a BCA Protein Assay Kit (Thermo; Cat#23227). 6.8 µg of each sample was reduced with DTT (final concentration 10mM) (Sigma; Cat#D0632) for 1 h at room temperature. Afterwards, iodoacetamide (IAA) (Sigma; Cat#I6125) was added to a final concentration of 50mM. The samples were incubated at room temperature for 1 h in the dark, with the reaction being stopped by addition of 10mM DTT. After precipitation of proteins using methanol/chloroform, the semi-dry protein pellet was dissolved in 25µL of 8 M urea in 20mM EPPS (pH 8.5) (Sigma; Cat#E9502) and was then diluted with EPPS buffer to reduce urea concentration to 4 M. Lysyl endopeptidase (LysC) (Wako; Cat#125-05061) was added at a 1:75 w/w ratio to protein and incubated at room temperature overnight. After diluting urea to 1 M, trypsin (Promega; Cat#V5111) was added at the ratio of 1:75 w/w and the samples were incubated for 6 h at room temperature. Acetonitrile (Fisher Scientific; Cat#1079-9704) was added to a final concentration of 20% v/v. TMTpro reagents (Thermo; Cat#90110) were added 4x by weight to each sample, followed by incubation for 2 h at room temperature. The reaction was quenched by addition of 0.5% hydroxylamine (Thermo Fisher; Cat#90115). Samples were combined, acidified by trifluoroacetic acid (TFA; Sigma; Cat#302031-M), cleaned using Sep-Pak (Waters; Cat#WAT054960) and dried using a DNA 120 SpeedVac™ concentrator (Thermo). Samples were then resuspended in 20mM ammonium hydroxide and separated into 96 fractions on an XBrigde BEH C18 2.1 × 150 mm column (Waters; Cat#186003023), using a Dionex Ultimate 3000 2DLC system (Thermo Scientific) over a 48 min gradient of 1–63%B (*B* = 20 mM ammonium hydroxide in acetonitrile) in three steps (1–23.5%B in 42 min, 23.5–54%B in 4 min and then 54–63%B in 2 min) at 200 µL min^−1^ flow. Fractions were then concatenated into 24 samples in sequential order (e.g. 1, 25, 49, 73). After drying and resuspension in 0.1% formic acid (FA) (Fisher Scientific), each fraction was analyzed with a 90 min gradient in random order.

### LC-MS analysis

Samples were loaded with buffer A (0.1% FA in water) onto a 50 cm EASY-Spray column (75 µm internal diameter, packed with PepMap C18, 2 µm beads, 100Å pore size) connected to a nanoflow Dionex UltiMate 3000 UPLC system (Thermo) and eluted in an increasing organic solvent gradient from 4 to 28% (B: 98% ACN, 0.1% FA, 2% H_2_O) at a flow rate of 300nLmin^−1^. Mass spectra were acquired with an orbitrap Fusion Lumos mass spectrometer (Thermo) in the data-dependent mode with MS1 scan at 120,000 resolution, and MS2 at 50,000 (@200 *m/z*), in the mass range from 400 to 1600 *m/z*. Peptide fragmentation was performed via higher-energy collision dissociation (HCD) with energy set at 35 NCE.

### Protein identification and quantification

The raw data from LC-MS were analyzed by MaxQuant, version 1.6.2.3 [74]. The Andromeda engine [75] searched MS/MS data against UniProt complete proteome database (*Mus musculus*, version UP000000589, 22,137 entries). Cysteine carbamidomethylation was used as a fixed modification, while methionine oxidation and protein N-terminal acetylation were selected as a variable modification. Trypsin/P was selected as enzyme specificity. No more than two missed cleavages were allowed. A 1% false discovery rate was used as a filter at both protein and peptide levels. First search tolerance was 20ppm (default) and main search tolerance was 4.5 ppm (default), and the minimum peptide length was 7 residues. After removing all the contaminants, only proteins with at least two unique peptides were included in the final dataset. Protein abundances were normalized by the total protein abundance in each sample in deep datasets. In the original dataset, protein abundances were normalized to ensure same median abundance across all channels in all replicates. Then for each protein log2-transformed fold-changes were calculated as a log2-ratio of the intensity to the median of all control replicates. All the proteins quantified in each experiment were used as the background.

## Statistical analysis

Statistical analysis was performed using GraphPad Prism v6.0. Statistical comparison of two unpaired groups was carried out using Mann-Whitney U non-parametric test unless stated otherwise. Welch’s variant of the Student t-test was used in indicated experiments. P-values under 0.05 were considered statistically significant and are denoted as * (p<0.05) or **(p<0.01).

## Data availability

The mass spectrometry data that support the findings of this study have been deposited in ProteomeXchange Consortium (https://www.ebi.ac.uk/pride/) via the PRIDE partner repository [76] with the dataset identifiers PXD025319.

## Acknowledgements

This work was supported by grants from the Knut and Alice Wallenberg foundation, the Swedish Medical Research Council, the Swedish Foundation for Strategic Research, the Hungarian Thematic Excellence Programme (TKP2020-NKA-26), and AstraZeneca. We thank F. Fiore for the construction of the *Ptpn22*^C129S^ mice. The work performed at Centre d’Immunophénomique was supported in part by the Investissement d’Avenir program PHENOMIN (ANR-10-INBS-07, to B.M.).

## Author contributions

JJ planned and performed data involving the mouse work, analyzed the data and wrote the first draft of the manuscript. B.M. conceived and developed the *Ptpn22*^C129S^ mice. FF and CMH contributed to the initial steps of the project. YC performed the analysis of the recombinant PTPN22 molecules. QC, MD and EA designed, planned and supervised the work and analysis of the recombinant PTPN22 proteins and EA made critical contributions to the writing of the manuscript. AAS, MG, and RZ contributed with mass spectrometry data. AÅ and RM contributed to the planning and supervision of the project. All authors revised and approved the manuscript. R.H. planned and supervised the project and takes the overall responsibility.

## Supplementary Figures

**Fig. S1.**
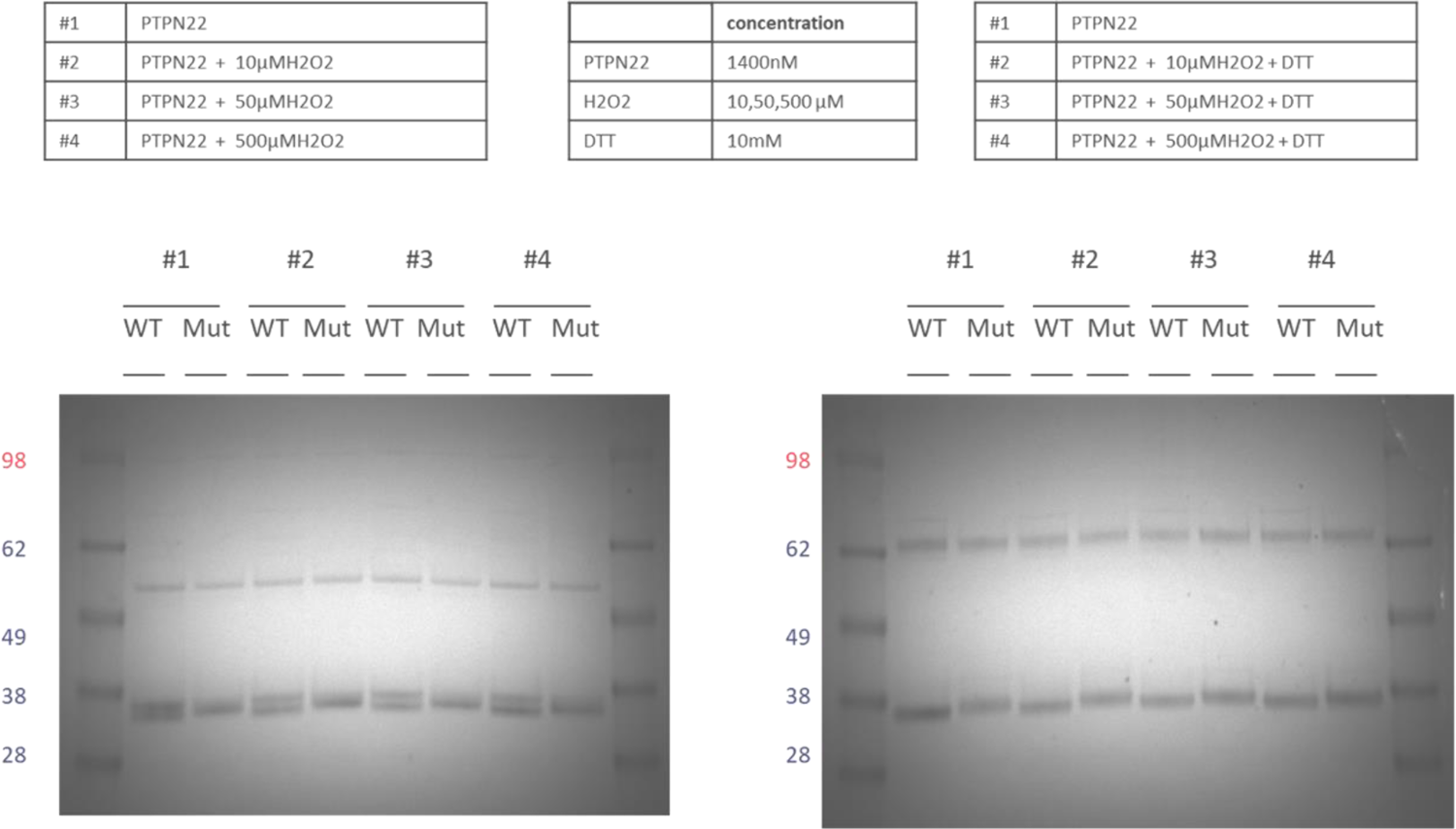
Reproducibility and H_2_O_2_-concentration dependence of the double band appearing in oxidized PTPN22 but not in PTPN22^C129S^, always fully reducible using subsequent treatment with DTT.

**Fig.S2.**
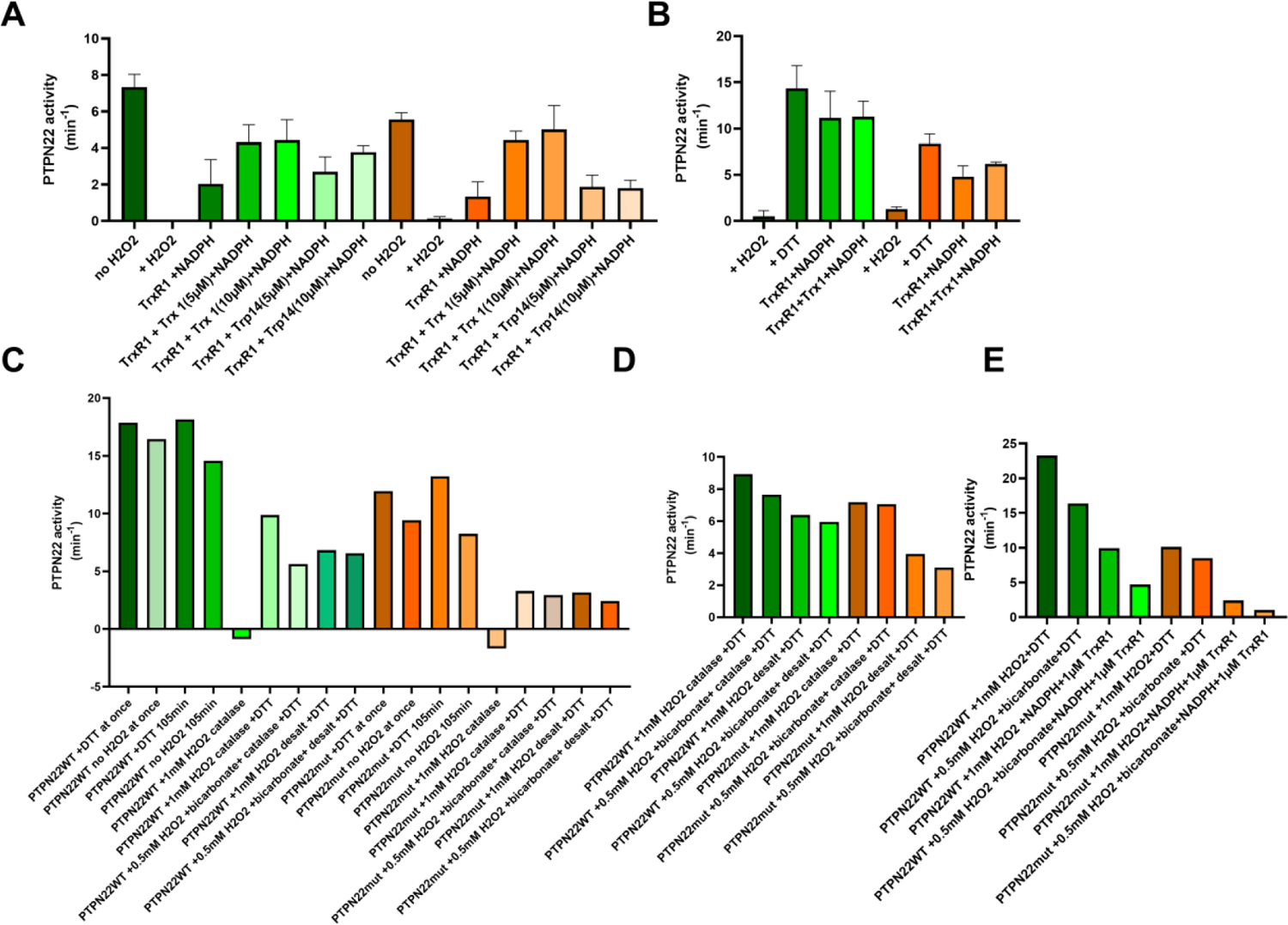
C129S mutant PTPN22 is less sensitive to TrxR1 reactivation as compared to WT PTPN22. **A)** Reduced wildtype (green) and C129S mutant (orange) PTPN22 (1400 nM) was pre-oxidized by H_2_O_2_ and then reactivated with TrxR1 (0.05 μM) (18 units/mg) and NADPH (300 μM) with or without TRP14 (5 or 10 μM) or Trx1 (5 or 10 μM), as indicated. After incubation for 60 min at 37°C, samples were analyzed for PTP activity. **B)** Reduced wildtype (green) and C129S mutant (orange) PTPN22 (1400nM) was treated with H_2_O_2_ (100 μM) for 30 min and then desalted to remove residual H_2_O_2_ with subsequent reactivation with DTT (10 mM) or TrxR1 (2.5 μM) (18 units/mg) with or without NADPH (300 μM) and Trx1 (10 μM). After 60 min at 37 °C, samples were analyzed for PTP activity. **C)** Reduced wildtype (green) and C129S mutant (orange) PTPN22 (1400nM) was oxidized using H_2_O_2_ with or without addition of 25 mM bicarbonate as described in the main text, but some groups included catalase and some reactivated with 10 mM DTT, as indicated. After 60 min at 37 °C, samples were analyzed for PTP activity. **D-E)** Reduced wildtype (green) and C129S mutant (orange) PTPN22 (1400 nM) was treated as described in C and reactivated with either DTT (10 mM) or TrxR1 (1μM) (18 units/mg) and NADPH (300μM), as indicated. After 60 min at 37 °C, samples were analyzed for PTP activity.

**Fig.S3:**
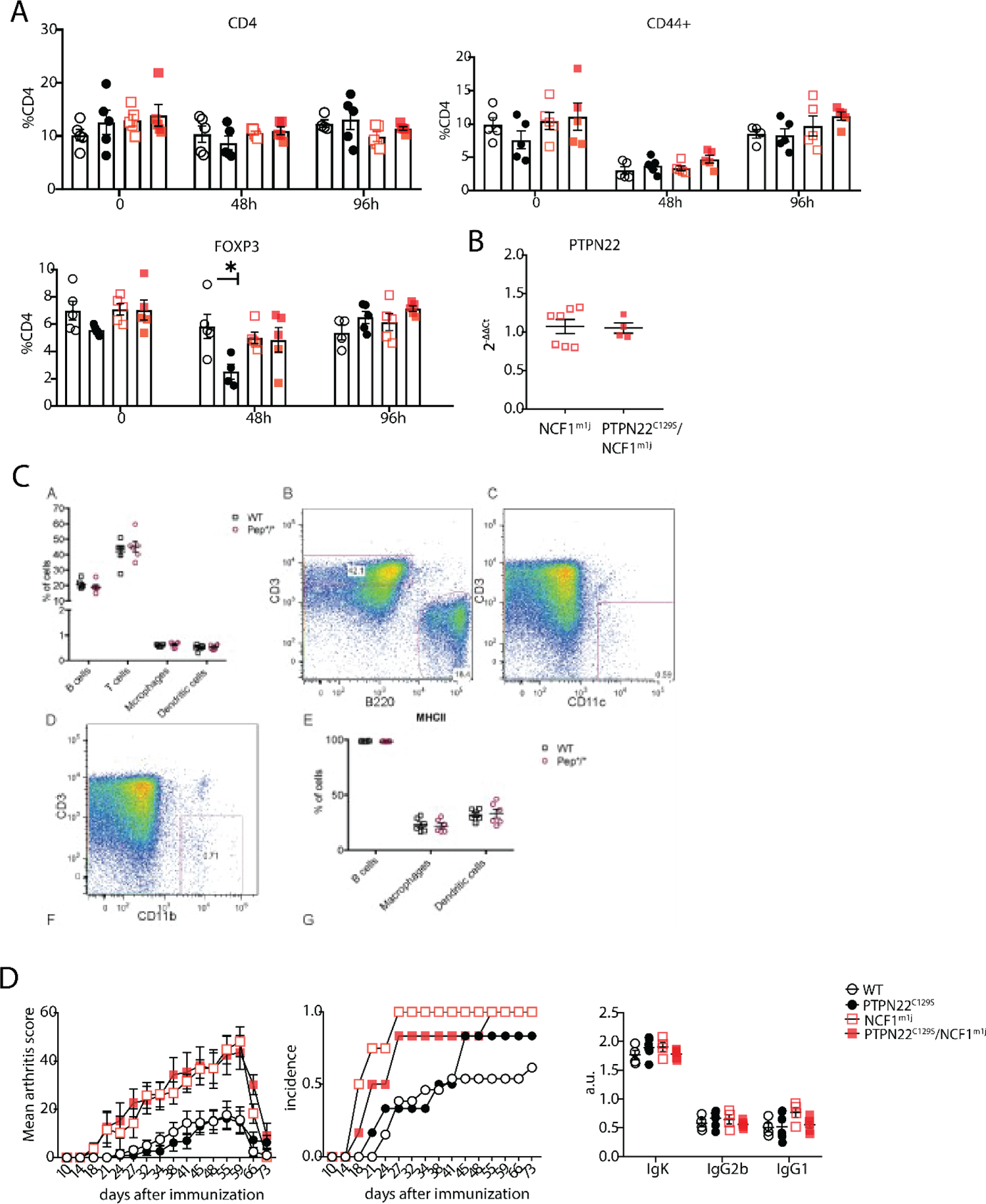
**A)** Flow cytometry measurement of indicated cell subsets in blood in mice 0, 48 and 96 hours after primary immunization with CII. **B)** Gene expression of PTPN22 in splenocytes from NCF1^m1J^ and PTPN22^C129S^/NCF1^m1J^ mice shown as fold change over NCF1^m1J^. **C)** Percentage of B cells (B220+), T cells (CD3+), macrophages (CD11B+), dendritic cells (CD11B+) in peripheral lymphoid organs as well as MHC II expression on B cells, macrophages and dendritic cells. **D)** Clinical score (mean±SEM), incidence and serum antibody levels at termination of collagen induced arthritis in littermate mice.

**Supplemental Table 1:**
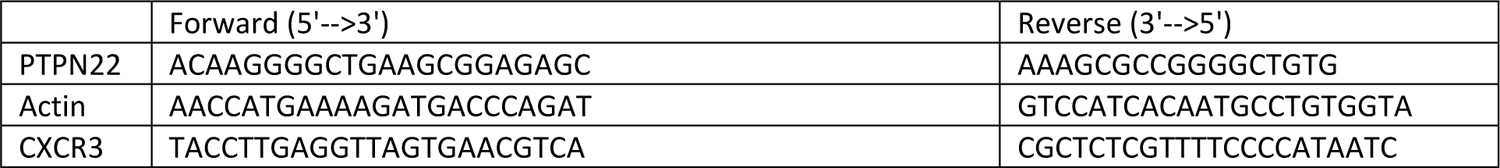
Primer list.

**Supplemental Table 2:**
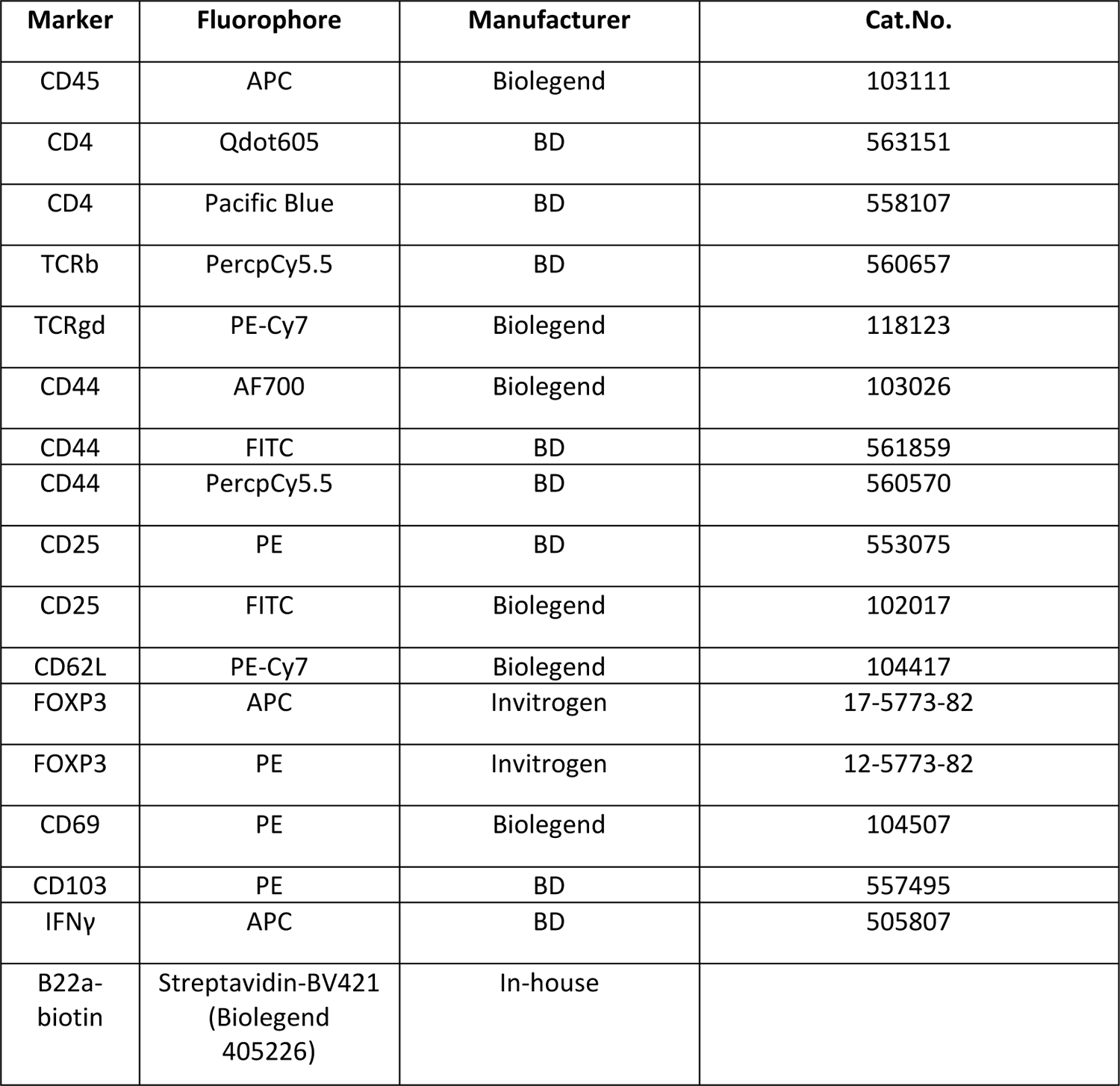
**antibody list**

| Marker | Fluorophore | Manufacturer | Cat.No. |
| --- | --- | --- | --- |
| CD45 | APC | Biolegend | 103111 |
| CD4 | Qdot605 | BD | 563151 |
| CD4 | Pacific Blue | BD | 558107 |
| TCRb | PercpCy5.5 | BD | 560657 |
| TCRgd | PE-Cy7 | Biolegend | 118123 |
| CD44 | AF700 | Biolegend | 103026 |
| CD44 | FITC | BD | 561859 |
| CD44 | PercpCy5.5 | BD | 560570 |
| CD25 | PE | BD | 553075 |
| CD25 | FITC | Biolegend | 102017 |
| CD62L | PE-Cy7 | Biolegend | 104417 |
| FOXP3 | APC | Invitrogen | 17-5773-82 |
| FOXP3 | PE | Invitrogen | 12-5773-82 |
| CD69 | PE | Biolegend | 104507 |
| CD103 | PE | BD | 557495 |
| IFNγ | APC | BD | 505807 |
| B22a-biotin | Streptavidin-BV421<br>(Biolegend 405226) | In-house |  |

